# A 20S proteasome receptor for degradation of intrinsically disordered proteins

**DOI:** 10.1101/210898

**Authors:** Assaf Biran, Nadav Myers, Julia Adler, Karin Broennimann, Nina Reuven, Yosef Shaul

## Abstract

Degradation of intrinsically disordered proteins (IDPs) by the 20S proteasome, unlike ubiquitin-dependent 26S proteasomal degradation, does not require proteasomal targeting by polyubiquitin. However, how these proteins are recognized by the proteasome was unknown. We report here on a mechanism of 20S proteasome targeting. Analysis of protein interactome datasets revealed that the proteasome subunit PSMA3 interacts with many IDPs. By employing in vivo and cell-free experiments we demonstrated that the PSMA3 C-terminus binds p21, c-Fos and p53, all IDPs and 20S proteasome substrates. A 69 amino-acids long fragment is autonomously functional in interacting with IDP substrates. Remarkably, this fragment in isolation blocks the degradation of a large number of IDPs in vitro and increases the half-life of proteins in vivo. We propose a model whereby the PSMA3 C-terminal region plays a role of substrate receptor in the process of proteasomal degradation of many IDPs.

## Introduction

Protein degradation plays key roles in diverse cellular processes and cell fate determination including proliferation, differentiation, death, antigen processing, DNA repair, inflammation and stress response as reviewed ^1,2^. A major regulatory route of protein degradation is controlled by the proteasome particles. Failure of the proteasome system and the resulting changes in protein homeostasis has been linked with human diseases and pathologies ^2–4^.

The 26S proteasome is an abundant cellular complex catalyzing protein degradation. It contains a 20S barrel-shaped proteolytic core particle, capped at one or both ends by the 19S regulatory complexes. The 20S proteasome is composed of four stacked rings in a barrel shape, two PSMA and two PSMB rings. Proteolytic activity resides in the chamber formed by the inner PSMB rings. The outer PSMA rings are identical and each has seven distinct subunits. The N-termini of the PSMA subunits form a gated orifice controlling substrate entry into the proteasome (Figure 1a). The 20S proteasome regulatory particles, including 19S, PA28, PA200, each interact with the PSMA ring, modulating its activity by opening the narrow entrance into the orifice and improving accessibility of substrates into the catalytic chamber ^5–10^

Proteins destined for degradation are first identified as “legitimate” substrates by the proteasomes prior to undergoing degradation ^11,12^. To this end, protein substrates target the proteasome via protein-protein interaction. The major targeting mechanism is the ubiquitin-dependent pathway. Polyubiquitin chains are covalently attached to the substrates, marking them for proteasome recognition and subsequent degradation. The polyubiquitin chain binds the 19S regulatory particle of the 26S proteasome directly or through transiently-associated 19S proteins ^13–15^. To date, three of the subunits of the 19S particle, namely Rpn1, Rpn10 and Rpn13 were identified as ubiquitin receptors ^16^. In the second pathway, no major prior protein modifications are required for targeting to the proteasomes. A well-known example of this pathway is ODC and antizyme. Binding of ODC by antizyme leads to proteasomal association and degradation ^17,18^. However, a growing number of proteins undergo proteasomal degradation by other mechanisms ^19–25^. Since interaction of the substrate with the proteasome is a requisite step in proteasomal degradation, the question of how these substrates are targeted to the proteasomes remained open. A considerable number of proteins undergo ubiquitin-independent degradation and by large are either completely or partially intrinsically disordered ^26^.

**Figure 1:**
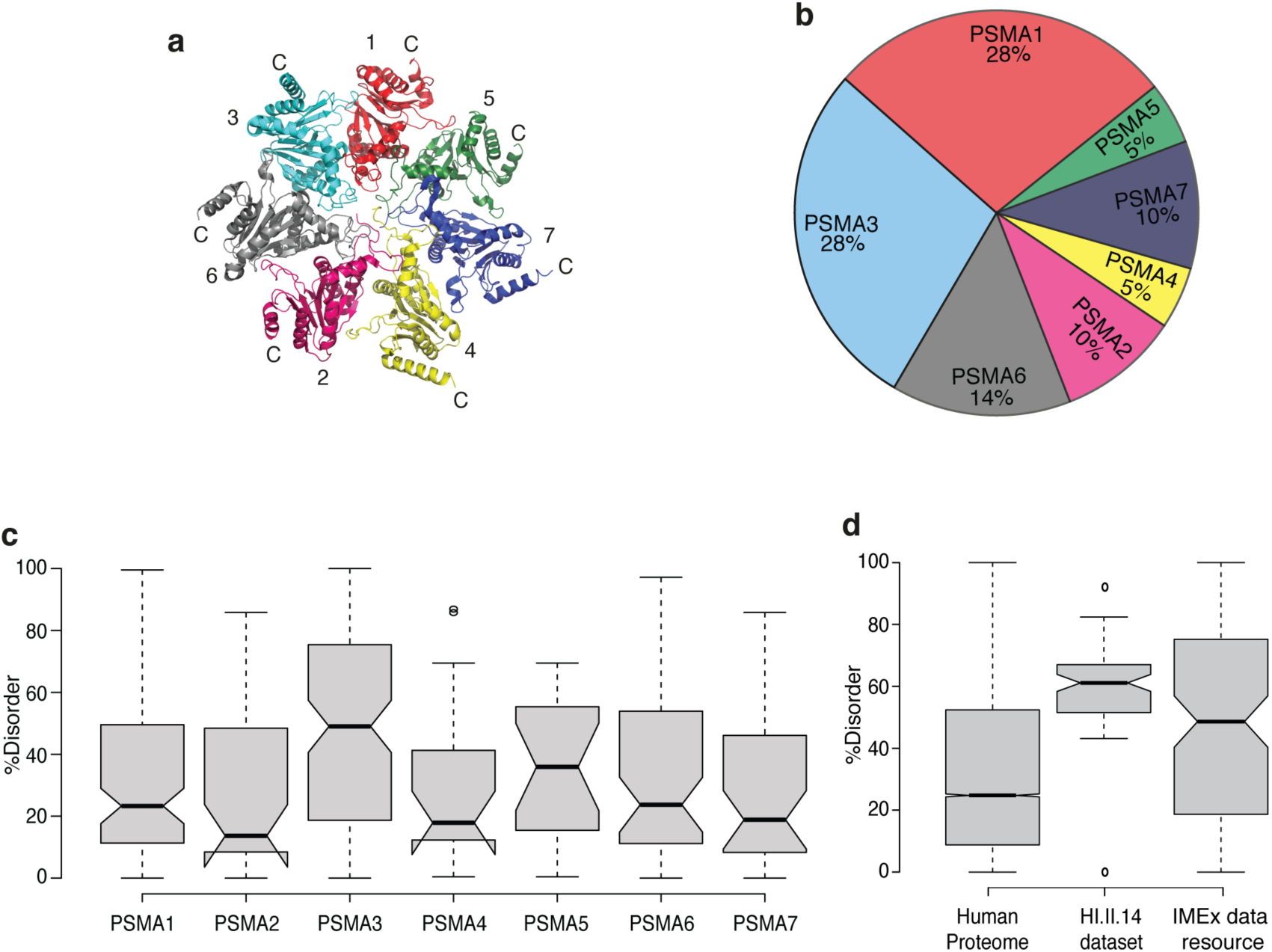
PSMA3 is a protein interaction hub of IDPs. (a) Crystal structure of PSMA ring adapted from Schrader *et al.*, 2016. PSMA subunits are identified by numbers. The N termini of the PSMA subunits protrude into the center of the ring, forming a gate restricting access into the 20S proteasome. (b) Pie chart presenting identified protein interactions of each PSMA subunit as a percentage of all identified protein interactions with PSMA subunits. We used the IMEx data resource to assemble an interaction list for the subunits. (c-d) Boxplot presenting the fraction of disordered residues found in the interacting proteins' sequences. Non-overlapping notches gives a 95% confidence that medians differ. Disordered residues were predicated with the IUPred algorithm. (c) Distribution of PSMA subunits interacting proteins from IMEx data resource. (d) Distribution of human proteome, PSMA3 interacting proteins from HI.II.14 dataset and IMEx data resource. See also supplementary table 1.

Over the last two decades many proteins have been identified as containing extensive disordered regions, and some proteins are even completely disordered under physiological conditions ^27,28^. These proteins were termed either as natively unfolded ^29^, intrinsically disordered proteins (IDPs) ^27,28^ and 4D proteins ^30^. IDPs are involved in many key cellular processes, including transcription regulation and signal transduction ^31^.

IDPs undergo 26S proteasomal degradation by the ubiquitination pathway. However, certain IDPs were shown to also undergo ubiquitin-independent proteasomal degradation in vitro using purified 20S proteasomes. The fact that these proteins are intrinsically unfolded negates the requirement of the 19S regulatory particle in unfolding the substrates. In fact, IDPs degradation by the 20S proteasome in vitro can be implemented in operational definition of this group of proteins ^30^. The notion is that this particle is gated and found in a latent state ^14^. However, allosteric mechanisms were suggested in opening the gate ^32^ and whether this is the case with the 20S particle is an open question.

Certain observations hint toward the possibility that the 20S proteasome is functional in the cells ^33^, especially under stringent conditions ^34–38^. Furthermore, in certain cases the physiological importance of the in vivo 20S activity was addressed ^39–43^. Since in this process ubiquitin is dispensable and the 19S particle-associated substrate receptors are absent, the mechanism of 20S particle targeting remained an open question.

We analyzed protein-protein interaction datasets and found that PSMA3, a 20S particle subunit, is an interaction hub for a subset of IDPs. We adopted the BiFC technique to validate our finding and mapped the interaction domain to the PSMA3 C-terminus. We further show that the PSMA3 C-terminal region works in heterologous contexts and in isolation to trap substrate candidates. Data obtained from *in vitro* and in cell experiments revealed that the trapper functions in facilitating IDPs ubiquitin-independent degradation. We propose a model whereby the 20S catalytic particle has a substrate receptor to trap certain IDPs for degradation.

## Results

### PSMA3 as an IDP-binding hub of the 20S proteasome

We have previously reported that certain intrinsically disordered proteins undergo proteasomal degradation by the 20S catalytic subunit ^39,41,42,44–46^. Given the fact that this process is ubiquitin-independent, the question of how the IDP substrates are recognized by the 20S proteasomal complex remained an open question. We assumed that an inherent component of the 20S complex might play a role of IDP trapper. Our assumption was that the putative IDP trapper is likely to be located in the PSMA ring as this ring forms the entry into the catalytic chamber (Figure 1a). To challenge this model we took advantage of the interactome data sets with the rationale that the PSMAs interacting proteins are potential 20S substrates. We analyzed the IMEx data resource which searches different databases of large-scale protein-protein interaction screens ^47^. The analysis revealed that PSMA3 and PSMA1 are the preferred protein-binding constituents (Figure 1b and supplementary table 1). Next, using the IUPred algorithm ^48^ we evaluated the percent disorder of the PSMAs interacting proteins. We found that the PSMA3-interacting proteins are uniquely highly enriched for IDPs (Figure 1c). We also compared PSMA3-interacting proteins found in the IMEx data resource to PSMA3-interacting proteins found in the HI.II.14 dataset from the human interactome project ^49^. The interacting proteins found in the HI.II.14 dataset are also enriched with IDPs (Figure 1d). These analyses support the possibility that PSMA3 might play a role of IDP substrate trapper in the process of ubiquitin-independent 20S proteasomal degradation.

### Chimeric PSMA3 produce BiFC with p21

We adopted the bimolecular fluorescence complementation (BiFC) assay ^50,51^ as a strategy to examine PSMA3 interaction with IDPs in cells. In this assay, the reporter fluorescent protein (FP) is split into two fragments; the C-terminus FPC and the N-terminus FPN, which upon their interaction emit a fluorescent signal. PSMA subunits were fused to FPC and the ubiquitin-independent proteasomal substrates were fused to FPN (Figure 2a). When PSMA and a potential substrate protein interact, the fluorescent fragments are brought into proximity, interact and fluorescence is restored (Figure 2b). The interacting chimeric subunits may emit a signal as free subunits or in the context of the proteasomes upon incorporation of the chimeric PSMA subunits.

**Figure 2:**
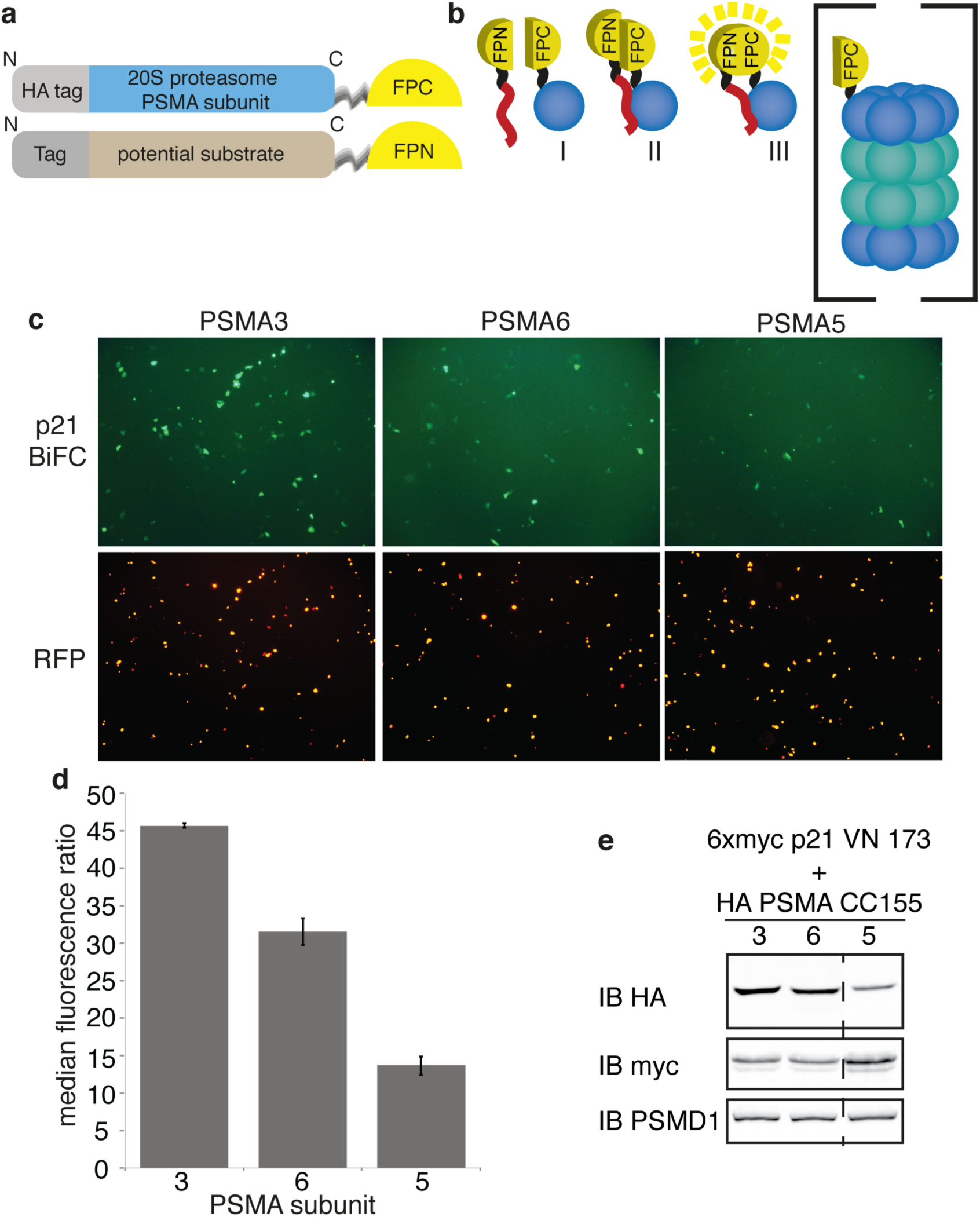
p21 interacts with PSMA3. (a) Illustration of constructs used for BiFC assay. A fluorescent protein was split to FPN (VN173) and FPC (CC155) terminal fragments. FPC is fused to a 20S proteasome PSMA subunit and FPN is fused to a potential substrate. (b) Schematic illustration of BiFC by three consecutive steps: Step I, the PSMA subunit fused to FPC and the substrate candidate fused to FPN are co-expressed; step II, the substrate interacts with the cognate PSMA subunit to increase substrate-proteasome accessibility; step III, the fluorescent protein refolds and fluorescence is restored. The chimeric PSMA subunit may or may not be incorporated into the 20S proteasome (c-e) HeLa cells were co-transfected with 6xmyc p21 FPN, chimeric PSMA3,5,6 subunits and H2B RFP. (c) Cells were examined for successful BiFC using a fluorescent microscope, 20x objective 48h post-transfection. (d) Intensities of at least 10,000 cells for each PSMA-p21 combination were recorded by flow cytometry. Standard deviation bars represent two independent experiments. (e) Expression level of the proteins in the cells presented in panel d was examined.

p21 is an IDP ^52,53^ which undergoes both ubiquitin-dependent and independent proteasomal degradation ^54,55^. In order to examine p21 interaction with PSMA3 we generated a chimeric 6xmyc p21 FPN. The myc tag minimizes p21 proteasomal degradation ^55^, to detect proteasomal recognition without compromising p21 level. BiFC signal of HeLa cells transiently co-transfected with PSMA3, 6, 5-FPC and p21-FPN was monitored by microscopy and FACS quantification. Chimeric PSMA 6 and 5-FPC were used as controls to evaluate the noise of the system. p21 gave the strongest signal when co-transfected with PSMA3 (figure 2c-d). We examined expression levels of the constructs in the cells used for FACS analysis to verify that the BiFC signal differences didn’t stem from different expression levels. PSMA3 and 6 were expressed to the same level thus excluding the possibility that BiFC efficiency differed because of expression (figure 2e). These data suggest that PSMA3 preferentially interacts with the intrinsically disordered protein p21.

### The PSMA3-FPC chimera is incorporated into proteasomes

In order to determine whether the PSMA3-FPC chimera is incorporated into the proteasomes we used native gel analysis. The analysis revealed that the PSMA3-FPC was incorporated into 20S and 26S proteasome complexes (Figure 3a). To quantify the fraction of the incorporated chimeric PSMA3 we conducted a successive proteasome depletion experiment (Figure 3b-c). The proteasomes were depleted from the cellular extract through immunoprecipitation of the endogenous 20S proteasome PSMA1 subunit and monitored for the presence of the chimeric PSMA3-FPC. We found that the PSMA3-FPC chimera was depleted as efficiently as the endogenous proteasome subunit PSMA1 (Figure 3d). Under this condition, and as expected, the 19S proteasome subunit PSMD1, was also depleted, although with lower efficiency (Figure 3d). These results suggest that the vast majority of the PSMA3-FPC chimeric protein is incorporated into the proteasomes.

**Figure 3:**
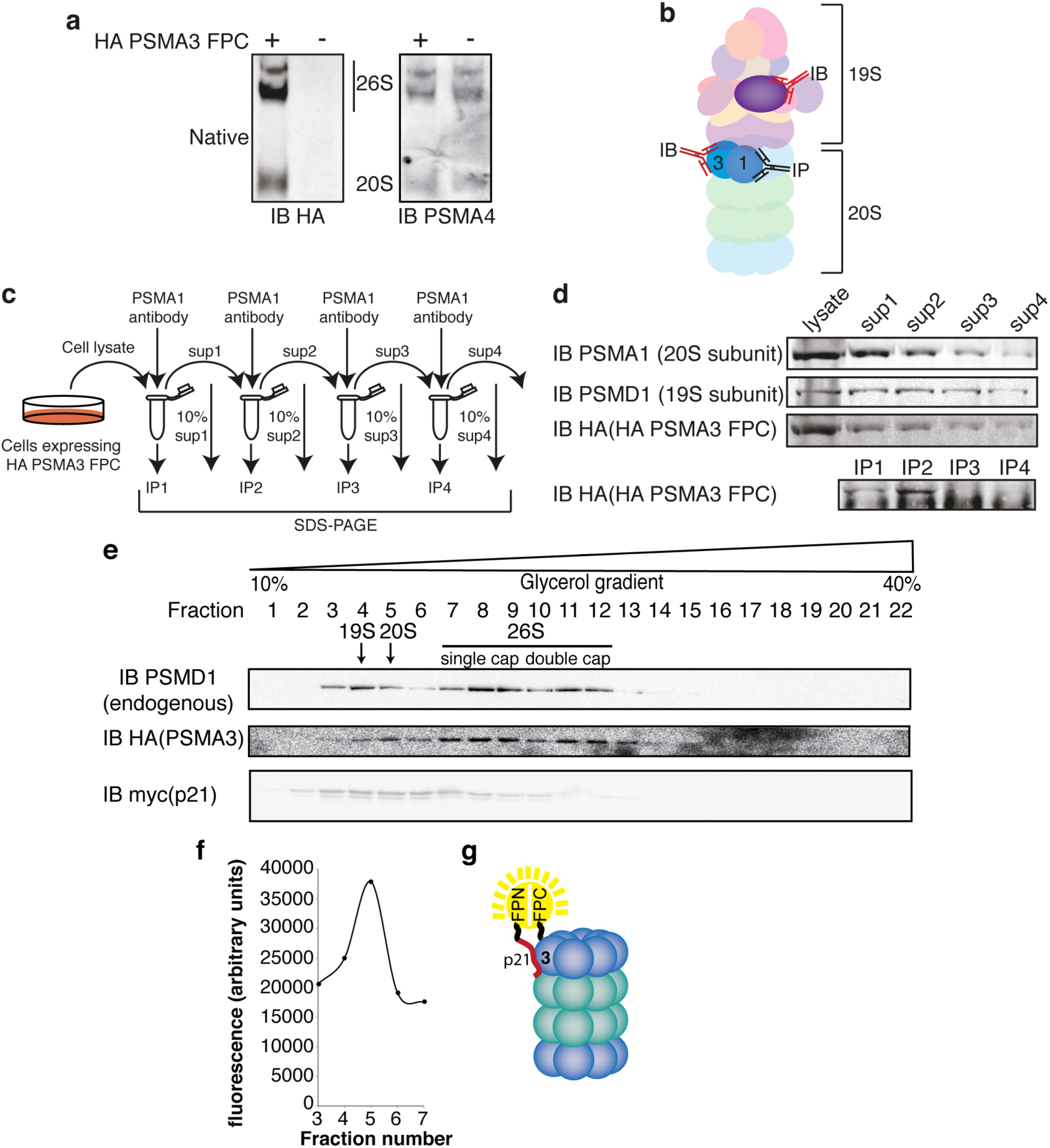
p21 FPN - PSMA3 FPC BiFC signal arises from interaction with intact 20S proteasome. (a) U2OS cells were transduced with PSMA3-FPC. Cell Lysates were enriched with proteasomes by ultracentrifugation and analyzed by native gel and immunoblot. Membrane was probed with an anti-HA antibody to detect the chimeric PSMA3 subunit, and with an antibody against the endogenous subunit PSMA4. (b) Schematic description of the co-immunoprecipitation steps to examine incorporation of chimeric PSMA3 subunit into proteasomes. The endogenous PSMA1 subunit was first immunoprecipitated and the level of the co-immunoprecipitated subunits was monitored using antibodies to detect the endogenous PSMD1, a subunit of the 19S proteasome and anti-HA to detect the chimeric PSMA3. (c) The schematic description of the experimental strategy of serial consecutive immunoprecipitation steps. (d) HEK293 cells expressing HA PSMA3 FPC were harvested 24h post transfection. Cells lysate was subjected to four subsequent immunoprecipitations of proteasomes via the endogenous PSMA1 subunit. Ten percent of cell lysate was kept for analysis after each immunoprecipitation. (e) HEK293 cells were transiently transfected with HA PSMA3 FPC and 6xmyc p21 FPN. Cells were harvested 48h post transfection, lysed, loaded on 11ml 10%-40% linear glycerol gradient and 0.5ml fractions were collected. Fractions were probed for the presence of HA PSMA3 FPC, 6xmyc p21 FPN and PSMD1, an endogenous 19S subunit. The PSMD1 fractionation profile enables determination of 20S and 26S (single and double cap) containing fractions. (f) BiFC signal of 20S proteasome containing fractions was quantified by a fluorometer. (g) Schematic summary model of this set of experiments.

### p21-FPN interacts with the proteasomal PSMA3-FPC

To address the question of the IDP interaction with the proteasomal PSMA3 we transfected the cells with the chimeric proteins and separated the complexes by glycerol gradient. We achieved clear separation of the distinct proteasome complexes from the whole cell extract. By immunoblotting with an antibody against PSMD1, a 19S subunit, we identified the fractions positive for the 19S (peaked at fraction 4), the single capped 26S and the double capped 26S particles (Figure 3e). The PSMA3-FPC protein was detected at the 20S region (peaked at fraction 5) and the two 26S complexes, consistent with the native gel data. The transfected p21-FPN was detected at a number of fractions, some of which contain the 20S peak of PSMA3 (fraction 5). We monitored GFP in the fractions and remarkably, the GFP level peaked at fraction 5 where the 20S PSMA3 peaked (Figure 3f). These data provide strong evidence for interaction of the chimeric p21-FPN with the PSMA3-FPC in the context of the whole 20S proteasome (Figure 3g).

### The PSMA3 C-terminus is sufficient to interact with p21

The PSMA subunits mainly differ at their C-termini ^56^ and PSMA3 C-terminus (Ct 187-255) is exposed enough in the 20S and 26S proteasomes to interact with IDP substrates (Figure 4a and supplement figure 1). Thus, we speculated that the PSMA3-Ct is the most likely p21-interacting region. To examine this possibility we constructed truncation mutants in the C-terminus of the chimeric PSMA3 (supplement figure 2a). Based on the secondary structure of PSMA3, the truncation was done at the flexible regions. Truncation longer than the last C-terminal 11 amino acid residues (Ct-Δ11) reduced the expression level of the subunit (supplement figure 2b). However, the Ct-Δ26 and Ct-Δ69 mutants were expressed to the same level yet the BiFC signal was markedly reduced in the Ct-Δ69 mutant (supplement figure 2c), suggesting that the PSMA3-Ct187-229 region recognizes p21. The crystal structure of PSMA3 suggests that the Ct187-229 region is adequately accessible to the surrounding proteins (Figure 4b).

**Figure 4:**
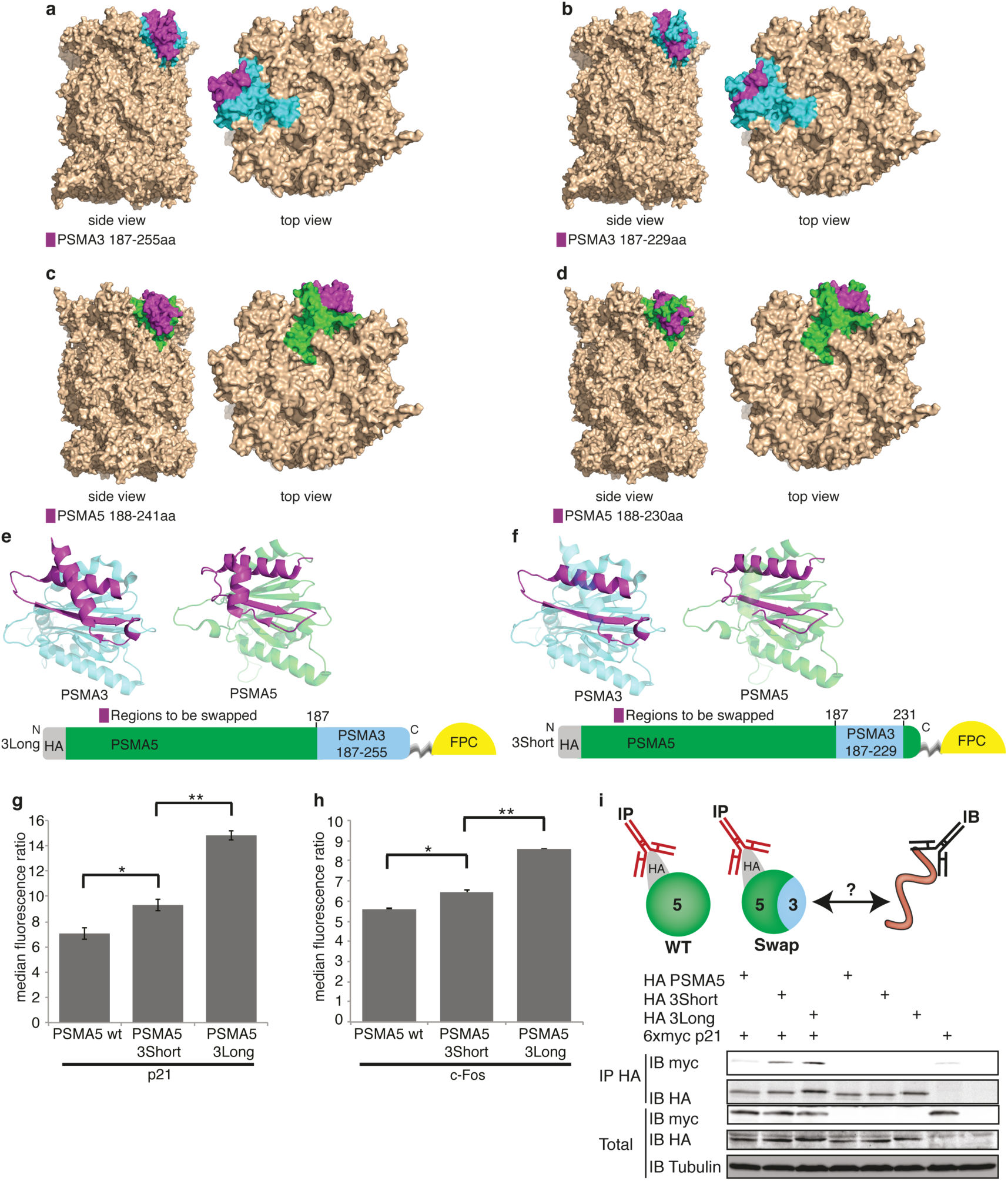
p21 and c-Fos interact with chimeric PSMA5 harboring PSMA3 C terminus. (a-d) crystal structure of the 20S proteasome marking (a-b) PSMA3 Ct and (c-d) PSMA5 Ct. The indicated C-terminal portions are labeled in magenta, and the remainder of the PSMA3 and PSMA5 subunits are labeled in cyan and green, respectively. The structure was taken from Schrader *et al.*, 2016. (e-f) Illustration of mutant PSMA5 constructs used in our experiments. Amino acid notations refer to PSMA5 WT. Crystal structures adapted from Schrader *et al.*, 2016. (g-h) HeLa cells were transiently transfected with chimeric PSMA5 subunit, swap mutants, H2B RFP, 6xmyc p21 FPN and flag c-Fos FPN as indicated. (g) Fluorescence intensities of at least 10,000 cells for each PSMA5-p21 combination were recorded by flow cytometry. Standard deviation bars represent three independent experiments. *p-value=0.03 **p-value=0.001 using 2 sided student t test. (h) Fluorescence intensities of at least 8,500 cells for each PSMA5-Fos combination were recorded by flow cytometry. Standard deviation bars represent three independent experiments. *p-value=0.01 **p-value=0.003 using 2 sided student t test. (i) Upper panel, illustration of experimental methodology. Lower panel, HEK293 cells were transiently transfected as indicated with 6xmyc p21 and chimeric PSMA5 subunit. Cells were harvested 48 hours post-transfection, lysed and subjected to immunoprecipitation (IP) with HA beads to immunoprecipitate chimeric PSMA5 subunit. Total lysate and IP samples were analyzed by SDS-PAGE and immunoblotting.

We next asked whether this region is sufficient to interact with p21 by conducting fragment swapping experiments. We chose PSMA5 to be swapped with the PSMA3-Ct as the BiFC signal with PSMA5 was weak with minimal background (Figure 2d) and since the PSMA5-Ct faces outward (Figure 4c-d). Two chimeric PSMA5 ΔCt-3Ct were constructed; one with a long PSMA3 Ct fragment (Ct187-255) and the other with a shorter Ct187-229 fragment (Figure 4e-f). We also generated reciprocal PSMA3ΔCt-5Ct constructs (supplement figure 3a-b). However, these constructs were expressed at much lower levels than the control, preventing reliable analysis, and therefore were not subjected to further studies (supplement figure 3c). The PSMA5 ΔCt -3Ct both long and short versions were efficiently expressed (supplement figure 3d). FACS quantification showed that both PSMA5 ΔCt-3Ct long and short yielded better BiFC with p21 than PSMA5 wt (Figure 4g).

We next examined if c-Fos, another IDP ^57^, which also can undergo ubiquitin-independent degradation ^39,40^ interacts with PSMA3-Ct. PSMA5 ΔCt -3Ct long and short gave a higher BiFC with c-Fos than PSMA5 wt (Figure 4h). Notably, the longer PSMA5 ΔCt-3Ct construct was significantly more efficient in emitting fluorogenic signals with p21 and c-Fos (Figure 4g-h).

To further validate the role of PSMA3-Ct region in interacting with IDPs, we conducted co-immunoprecipitation experiments. For this set of experiments we used a 6xmyc-tagged p21 construct lacking the FPN moiety and found it to be co-immunoprecipitated with the chimeric PSMA5 ΔCt-3Ct long and short swapped constructs but not with the naïve PSMA5 (Figure 4i). These data suggest that the PSMA3-Ct is sufficient in interacting with p21 also in the PSMA5 context.

### Recombinant PSMA3 trapper interacts with a subset of IDPs

To demonstrate that PSMA3-Ct plays a role of IDP trapper we examined its capacity to interact with IDPs in isolation using a recombinant GST fusion protein containing the putative PSMA3-Ct187-255 IDPs trapper. We used two controls, the GST-PSMA5-Ct chimeric protein, where the PSMA5-Ct188-241, which is structurally analogous to the trapper region of PSMA3 were fused to GST, and as a second control we used the recombinant GST (Figure 5a). HEK293 cell extract overexpressing 6xmyc p21 was incubated with purified GST-fusion proteins bound to glutathione—agarose beads and eluted fractions were analyzed by immunoblotting. Remarkably, the 6xmyc p21 was efficiently pulled down only with the GST-PSMA3-Ct putative trapper (Figure 5b).

**Figure 5:**
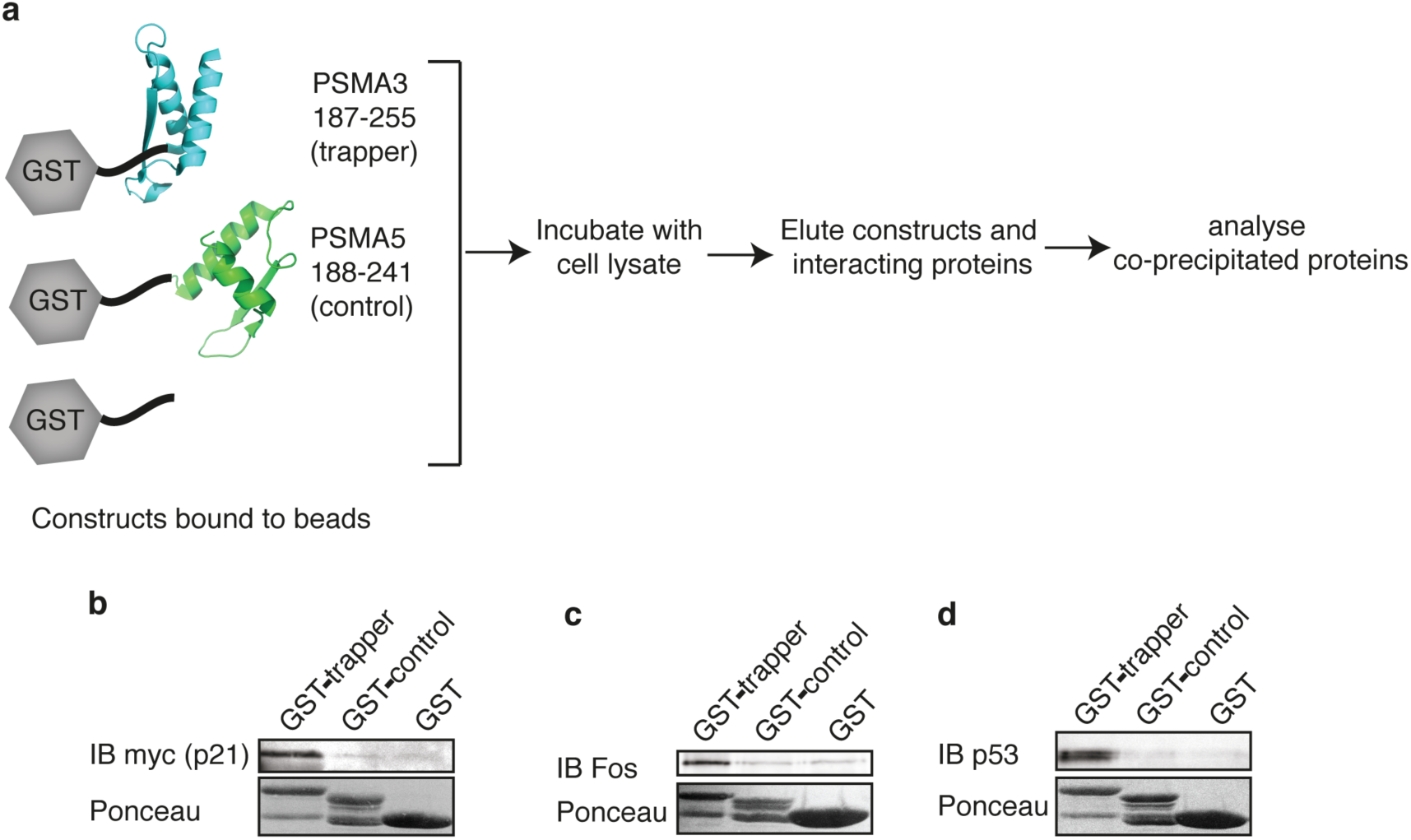
Isolated PSMA3 C-terminus interacts with three intrinsically disordered proteins. (a) Illustration of constructs used and experimental strategy. (b-d) Purified GST, GST PSMA3 trapper and GST PSMA5 C terminus bound to Glutathione agarose beads were incubated with HEK293 cell lysate overexpressing 6xmyc p21 (b) or naïve HEK293 cell lysate (c-d). GST constructs and the interacting proteins were eluted with 10mM reduced glutathione. GST constructs were visualized with Ponceau and interacting proteins by immunoblot (IB).

Next, we examined the recombinant trapper ability for pulling down specific endogenous IDPs, such as c-Fos and p53, which were shown to undergo ubiquitin-independent degradation ^41,42,58^. Remarkably, the PSMA3 trapper fragment specifically pulled down c-Fos and p53 (Figures 5c-d). The interaction was highly specific since neither GST nor GST-PSMA5-Ct ligands were active. These data suggest that the PSMA3-Ct region is an autonomously functioning trapper of certain IDPs.

### The PSMA3 trapper regulates p53 degradation

Two models of ubiquitin-independent degradation of IDPs are proposed (Figure 6a). Model I is a one step process, an IDP encounters the 20S proteasome and undergoes degradation. Model II is a two-step process, PSMA3 acts as a trapping mechanism for substrate-proteasome interaction to facilitate degradation. If model II is the correct one, an excess of the recombinant PSMA3-Ct trapper is expected to protect IDPs from ubiquitin-independent degradation in vitro. To test this possibility we conducted an in vitro reaction to degrade p53 by the 20S proteasome ^30,42^ We incubated purified recombinant human p53 with 20S proteasomes in the absence or presence of recombinant GST-proteins. Remarkably, addition of GST-PSMA3 trapper but not the control proteins significantly inhibited p53 degradation (Figure 6b-c). The purified GST fusion proteins do not inhibit 20S proteasome activity with the fluorogenic substrate Suc-LLVY-AMC (supplement figure 4), therefore, the process of inhibition of the p53 degradation is due to preventing interaction with PSMA3 trapper region.

**Figure 6:**
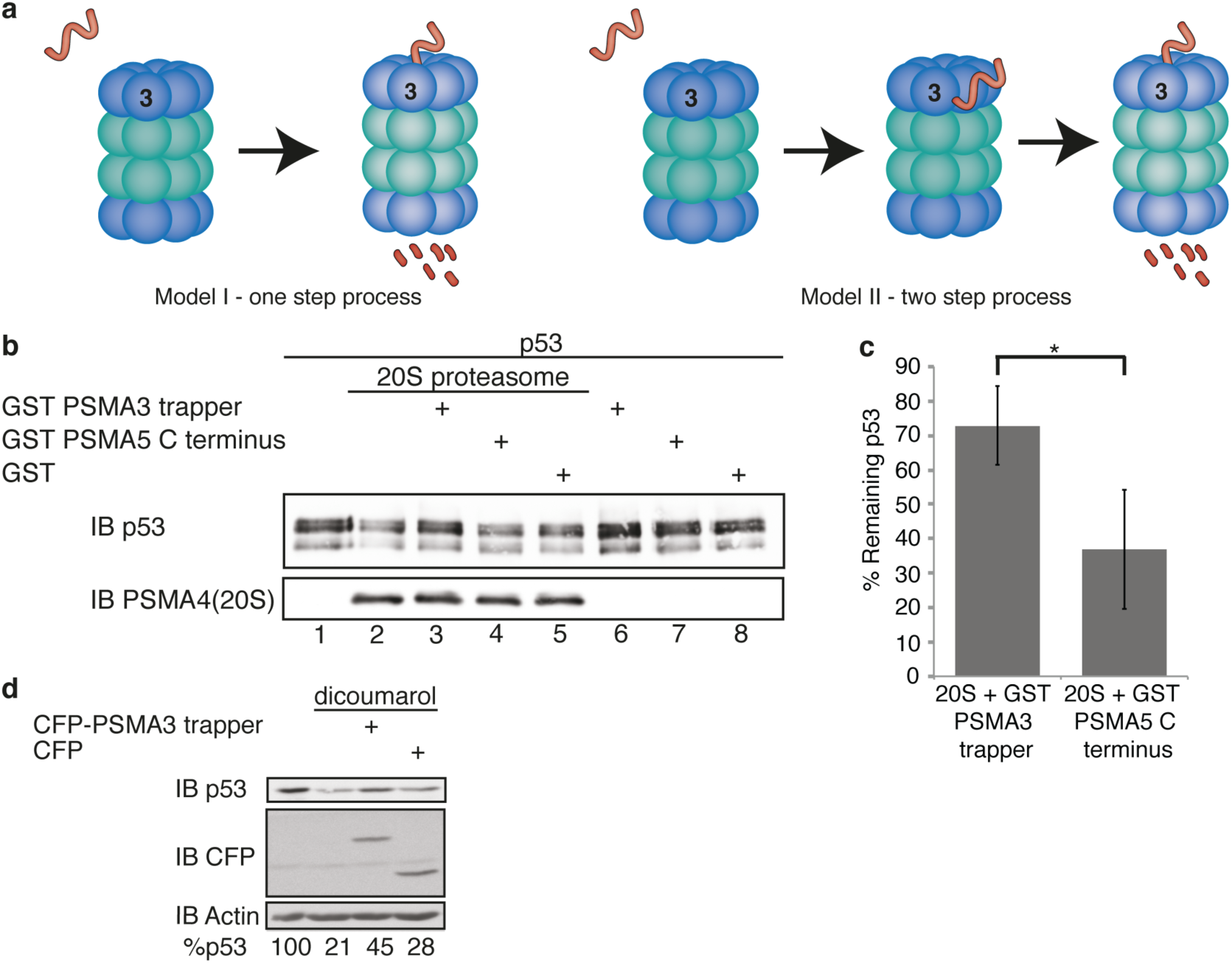
The PSMA3 substrate trapper inhibits p53 proteasomal degradation. (a) Illustration describing two possible models of ubiquitin independent degradation of IDPs. (b) p53 purified from recombinant baculovirus-infected cells was incubated 1h at 37^o^C with purified 20S proteasome in the presence or absence of the GST-PSMA3 trapper, GST-PSMA5 C-terminus and GST alone, as indicated. (c) Quantification of purified p53 degradation by 20S proteasome in the presence of GST-PSMA3 trapper or GST-PSMA5 C-terminus. Standard deviation bars represent four independent experiments. *p-value=0.02 using 2 sided student t test. (d) HCT116 cells were transfected with CFP-PSMA3 trapper and CFP as indicated. 24h post-transfection cells were incubated with or without 500μM dicoumarol for 5h. Cell extracts were analyzed by SDS-PAGE and immunoblotting (IB).

To examine the involvement of the PSMA3 trapper in p53 degradation in the cells we employed our published protocol ^59^. We have previously shown that NQO1 protects p53 from ubiquitin-independent proteasomal degradation and that dicoumarol, an NQO1 inhibitor, increases p53 proteasomal degradation (Figure 6d). Remarkably, this process was attenuated by over-expression of the CFP-PSMA3 trapper and not by the CFP control (Figure 6d). PSMA3 trapper does not change total ubiqitination level or structured protein degradation (Figure 6-figure supplement 2), ruling out the possibility of interfering with ubiquitin dependent degradation. These data suggest that the PSMA3 trapper fragment inhibits ubiquitin-independent p53 degradation both in vitro and in the cells.

### The PSMA3 trapper regulates c-Fos degradation

We next examined c-Fos degradation using an experimental setting where it undergoes ubiquitin-independent degradation in cells ^39,40^ (Figure 7a). The level of c-Fos accumulation after serum induction is modulated by ubiquitin-independent proteasomal degradation. We ectopically expressed CFP-PSMA3 trapper and CFP in this setting and examined the effects on c-Fos accumulation kinetics. c-Fos accumulation was higher in the presence of CFP-PSMA3 trapper and lasted for a longer period (Figure 7b). Furthermore, we found that the c-Fos half-life was increased in the presence of the CFP-PSMA3 trapper (Figure 7c). These results demonstrate that the PSMA3 trapper regulates c-Fos proteasomal degradation.

**Figure 7:**
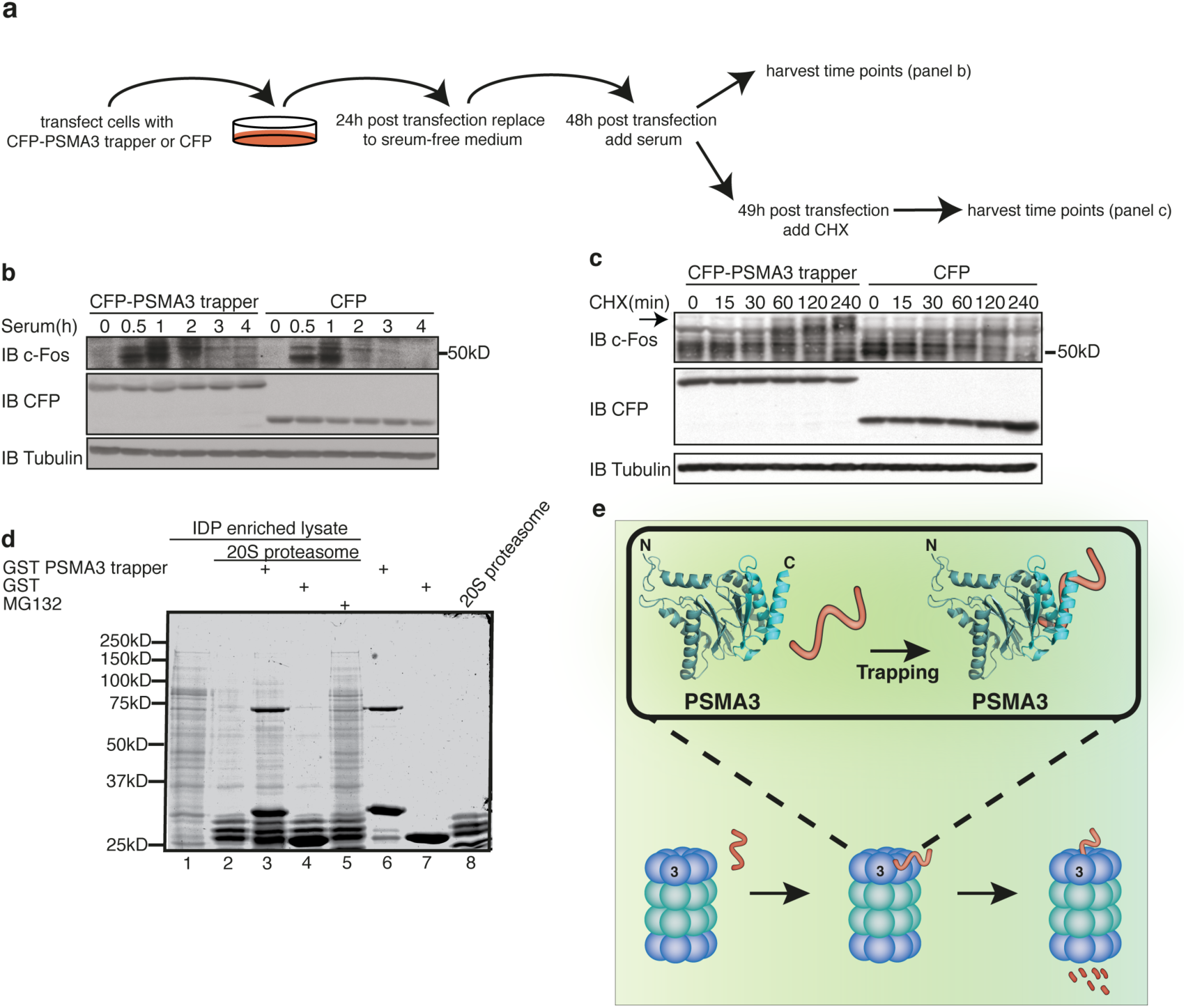
Isolated PSMA3 substrate trapper inhibits many IDPs’ 20S proteasomal degradation. (a) Illustration of experimental methodology. (b) Accumulation of endogenous c-Fos in response to serum induction was examined in the presence of CFP-PSMA3 trapper or CFP in HCT116 cells. Cells were transfected with the constructs as indicated. 24h post-transfection cells were serum starved for 24h. Serum was added and cells were harvested at the indicated time points. Immunoblot (IB). (c) Cells were treated as in panel b. 1h after serum induction 350μM cycloheximide was added for the specified time. Arrow marks slow mobility form of c-Fos. (d) HEK293 IDP enriched lysate was incubated 3h at 37°C with purified 20S proteasome, GST-PSMA3 trapper and GST as indicated. Proteins were visualized with InstantBlue stain. (e) Illustration describing our model of ubiquitin-independent degradation of IDPs.

### PSMA3 trapper facilitates 20S proteasomal degradation of many IDPs in vitro

We next examined the role of the PSMA3 trapping region in 20S proteasomal degradation of various IDPs. To this end we incubated an enriched IDPs lysate (Csizmók et al. 2006; Galea et al. 2006; Irar et al. 2006; Galea et al. 2009) with purified 20S proteasome. As expected ^30^ IDP enriched lysate was massively degraded by the 20S proteasome (Figure 7d lanes 1 and 2). Remarkably, when the recombinant GST-PSMA3 trapper was added, degradation was markedly compromised (Figure 7d lane 3). The decoy effect was specific and was not recapitulated by the control recombinant GST (Figure 7d lane 4). The lysate was prepared by heat treatment and therefore the protein complexes are expected to be dissociated, a critical step in reducing indirect trapper association and conducting *in vitro* 20S proteasomal degradation. These results suggest that the 20S proteasomal degradation of a large number of IDPs is regulated by the PSMA3 trapper.

## Discussion

### PSMA3 is an IDP-interacting hub of the proteasome

A key regulatory process in proteostasis is ubiquitin-dependent proteasomal degradation. In this process substrates are ubiquitinated and targeted to the 26S proteasome for degradation. Ubiquitinated substrates interact directly with the 19S subunits Rpn1, Rpn10 and Rpn13, as a mechanism of substrate recruitment ^16^. In this study, we examined the possibility of substrate recognition by an alternative and ubiquitin-independent mechanism. This study was motivated by the findings that certain proteins, in particular IDPs, are proteasomally degraded in the absence of ubiquitination ^26^. Based on the analysis of interactome data sets we identified the 20S proteasome subunit PSMA3 as an IDP interacting hub and assumed that PSMA3 plays a role of an IDP receptor. We took a number of in vivo and cell free experimental approaches to validate this finding and to demonstrate that PSMA3 interacts with and facilitates IDPs degradation. These findings led to the identification of a novel substrate trapper embedded in the C-terminus of PSMA3, a 20S proteasome subunit. This conclusion is based on the following observations. Using a BiFC system we demonstrated that the PSMA3 subunit and specifically its C-terminus interacts with the IDPs p21 and c-Fos. We further showed that the isolated PSMA3 C-terminus is sufficient to interact with p21, c-Fos and p53 using a co-immunoprecipitation strategy. Based on these attributes we termed the element a substrate trapper. Although we have directly investigated only the p21, p53 and c-Fos substrates, we assume that a larger number of IDPs are likely to be trapped. This view is based on our finding that excess of recombinant trapper, that was shown to be active in binding the tested proteins, markedly reduces the degradation of pools of cellular IDPs by the 20S proteasome.

### The proteasome species that degrade IDPs

A few species of proteasomes are found in cells and they differ by the type of regulatory particle associated with the catalytic 20S particle. Cells contain relatively large amounts of the 20S particles but whether they are physiologically functional is debatable. The accepted assumption is that this particle is in a latent state and tightly gated to minimize unscheduled protein degradation ^14^. However, given the fact that the mechanism of the gate opening neither in the context of the 26S proteasome nor the free 20S form is settled, we need to rely on empirical evidence for the possible in vivo activity of the 20S particle. For example, proteins damaged by oxidation were shown to undergo ubiquitin-independent degradation by the 20S proteasome ^33^. Both oxidative damage and prolonged cell starvation stress results in 26S proteasome disassembly and 20S accumulation ^34–36^. p53 and c-Fos are both stress response IDPs and both are 20S proteasome substrates ^39–42^. Furthermore, two independent groups have found that reducing the expression of the 19S regulatory particle subunits improves viability of cells treated with 20S proteasome inhibitors ^37,38^, attributing an active role to the 20S particle in cell fate determination. Finally, in the nervous system, membrane-associated 20S proteasomes modulate the calcium signaling induced by neuronal activity ^43^. All these findings support the possibility of at least a fraction of 20S particles playing physiological roles and that the identified PSMA3 IDP trapper is likely to regulate these processes.

Recently our lab reported the existence of another form of 26S proteasome. The 26S proteasome is also stable when it binds the small molecule NADH without the need for ATP ^64^. We have evidence that the 26S NADH proteasome can efficiently degrade IDPs, unlike 26S ATP-stabilized proteasome. In addition, the identified PSMA3 trapper region is exposed in the 26S proteasome (supplement figure 1). Thus, one can think of the possibility that the 26S NADH proteasome can trap IDPs through PSMA3.

### The PSMA3 substrate receptor

The PSMA subunits are highly similar but are unique at their C-terminus ^56^. The PSMA3 C-terminal region bears a helix-strand-helix structural motif (Fig 4). Deletion mutants revealed that the removal of the C-terminus α helix region, the last 26 aa residues, sharply reduces protein accumulation of this construct, therefore the trapper region determines PSMA3 protein level as well. The identified trapper region is rather charged, but since other PSMAs are also highly charged at their C-termini but lack IDP trapping activity, we assume that the charged residues might be required but are not sufficient to function as substrate trapper. All the PSMA subunits C-termini contain the helix-strand-helix motif but with a different helix-to-helix positioning, which is likely to contribute to the selection of the client proteins.

We demonstrated both *in-vitro* and in cells that the PSMA3 substrate trapper region is an autonomous module and in isolation is active in substrate binding. We took advantage of its autonomous behavior to sequester the client proteins in binding the 20S particle. The obtained data are consistent with the proposed model (Figure 7e) that trapper accessibility is a prerequisite step in substrate degradation. Mechanistically, the receptor-substrate interaction would position the substrate close to the 20S catalytic orifice. In addition, we would like to speculate that this interaction may facilitate orifice opening. In the 20S proteasome the orifice is occluded by the N termini of the PSMA ring subunits, thus creating a gate limiting access into the proteolytic chamber ^8,65,66^. Examination of the 20S proteasome by other methods revealed that the 20S gate can interchange between open and closed conformations ^67–69^. It is likely that the substrate-receptor interaction induces or stabilizes an open gate conformation enabling access into the catalytic chamber. This possibility is supported by reports showing that substrates of the 20S proteasome stimulate its activity ^70^ and an allosteric effect inducing an open gate conformation upon interaction with the peptidase active sites ^32^. Interestingly, along this line, it has been reported that artificial targeting of a substrate to PSMA3 in yeast stimulates degradation ^71^. Furthermore, the gate in the 26S proteasome is also largely found in a closed conformation ^9,10,72^. However, interaction of ubiquitinated substrates with the proteasome through the 19S particle allosterically induces their degradation ^73^ most likely by stabilizing the open gate conformation in the 26S proteasome. The possibility that the substrate trapper plays double roles in recruiting the client proteins and in activating the 20S proteasome is intriguing and demands further investigation.

### Regulation of the trapper-mediated substrate selection

A key question is how the IDP trapper/degradation is regulated to permit substrate discrimination. It is well demonstrated that IDPs undergo rather extensive post-translational modifications ^74–76^. These modifications, largely S/T phosphorylation, might very well increase or decrease trapper binding. Also, previously we reported on nanny proteins that interact with IDPs to escape their proteasomal degradation ^77^. Nanny proteins therefore might function to help the substrate in escaping the trapper. According to these models, trapper interaction takes place by default unless the system instructs otherwise. This model is different from the ubiquitination process where the substrate is stable unless marked by ubiquitin.

## Materials and methods

### Tissue culture

The cell lines used were: HEK293, U2OS, HCT116 and HeLa. Cells were grown in DMEM supplemented with 8% fetal bovine serum, 100 units/ml penicillin, and 100 μg/ml streptomycin and cultured at 37°C in a humidified incubator with 5.6% CO2.

### Plasmids, transfection and infection

Plasmids used: PCDNA3 CFP and PCDNA3 CFP PSMA3 187-255aa. PSMA subunits (Kindly provided by Prof. K. Tanaka, Tokyo Metropolitan Institute of Medical Science, Japan) were cloned into pBiFC-VN173, a gift from Prof. Chang-Deng Hu (Addgene plasmid no. 22010). 6xmyc p21 (Kindly provided by Prof. Chaim Kahana, Weizmann Institute of Science, Israel), c-Fos and p53 were cloned into pBiFC-CC155, a gift from Prof. Chang-Deng Hu (Addgene plasmid no. 22015). HEK293 cells were transfected by the calcium phosphate method. HeLa and HCT116 cells were transfected with jetPEI (Polyplus-transfection SA, Illkirch, France).

### Immunoblot analysis

Cells were lysed with NP40 buffer (20mM Tris-HCl pH7.5, 320mM sucrose, 5mM MgCl_2_, 1% NP40) supplemented with 1mM Dithiothreitol (DTT) and protease and phosphatase inhibitors (Sigma). Laemmli sample buffer (final concentration 2% SDS, 10% glycerol, 5% 2-mercaptoethanol, 0.01% bromphenol blue, 0.0625 M Tris-HCl pH6.8) was added to the samples, heated at 95°C for 3 minutes and loaded on a polyacrylamide-SDS gel. Proteins were transferred to cellulose nitrate 0.45 μm membranes. Antibodies: Mouse anti-HA, mouse anti-Flag, mouse anti-actin, mouse anti-tubulin and mouse anti-human p53 Pab1801 were purchased from Sigma. Mouse anti-myc was produced by the Weizmann Institute Antibody Unit. Rabbit anti-PSMD1, a subunit of the 19S proteasome, was purchased from Acris. Rabbit anti-PSMA4, a subunit of the 20S proteasome ^80^, was kindly provided by Prof. C. Kahana, Weizmann Institute of Science, Israel. Secondary antibodies were horseradish peroxidase-linked goat anti-rabbit and anti-mouse (Jackson ImmunoResearch). Signals were detected using the EZ-ECL kit (Biological Industries).

### Glycerol gradient

HEK293 cell were lysed in 0.5ml NP40 buffer and loaded on an 11ml linear 10%-40% glycerol gradient and centrifuged 16 hours at 28,400 rpm, using rotor SW 41TI. 0.5ml fractions were collected and analyzed by western blot and fluorometer.

### Co-immunoprecipitation

Samples were incubated with primary antibody 16h. Samples were washed 6 times with NP40 buffer. Bound and associated proteins were eluted with Laemmli sample buffer or HA peptide (Sigma) according to standard protocol.

### Nondenaturing PAGE

Samples were prepared and run as described ^42^.

### BiFC analysis

Cells were co-transfected with PSMA subunit-FPC, potential substrate-FPN and H2B-RFP. Cells with successful BiFC are colored green (VFP) and H2B-RFP colors cell nuclei in red. For flow cytometry analysis, cells were harvested 48h post transfection, washed and resuspended in PBS. Samples were analyzed with BD LSR II flow cytometer using FACSDiva software (BD Biosciences). VFP and RFP intensities of RFP positive cells were recorded. Values of VFP and RFP fluorescence intensities of each cell were extracted using FlowJo software (FlowJo, LLC). BiFC signal was normalized to RFP signal per cell. The BiFC/RFP median is used as the ratio distribution is skewed ^81^.

### Purification of the 20S proteasome

We used a modified protocol based on Beyette, Hubbell and Monaco, 2001. Mouse livers were homogenized in buffer containing 50mM Tris-HCl (pH 7.5), 150mM NaCl and 250 mM sucrose. After subsequent rounds of centrifugation (1750xg 15min, 34,500xg 15min, 100,000xg 1h) proteasomes were pelleted by centrifugation at 150,000g for 6h, and re-suspended in buffer containing 20mM Tris-HCl (pH 7.7), 150mM NaCl and 15% glycerol. Sample was loaded on a Hiprep Q (GE Healthcare) column and proteasomes were eluted from the column with a gradient of 0.2–0.5M NaCl over 10 column volumes (CVs), and 0.5–1M NaCl over 5 CVs, collecting 2-mL fractions. Pooled proteasome fractions were supplemented to 1.75M (NH_4_)_2_SO_4_ and contaminating remaining proteins were removed by pelleting at 15,000g for 30min. Supernatant was loaded on a HiTrap Phenyl HP column (GE Healthcare), washed with 10 CVs of buffer containing 20mM Tris-HCl (pH 8) and 2M (NH_4_)_2_SO_4_ and eluted by 4 CVs of 2M-1.2M (NH_4_)_2_SO_4_, followed by a gradient from 1.2M–0M (NH_4_)_2_SO_4_ over 20 CV, collecting 1 mL fractions. Glycerol was added immediately to each fraction to a final volume of 20%, following elution, in order to preserve proteasome activity. Eluted fractions were dialyzed against a buffer containing 50mM Tris-HCl (pH 7.5), 5mM MgCl_2_ and 20% glycerol. Purest fractions containing proteasomes were pooled and concentrated using a Millipore 100kDa-cut off concentrator. Protein concentration was estimated by Bradford Assay, purification purity was visualized on an SDS-PAGE gel by a Coomassie stain (InstantBlue Expedeon) and Proteasome activity was determined by the ability to hydrolyze the fluorogenic peptide suc-LLVY-AMC. Proteasomes were aliquoted, flash-frozen in liquid nitrogen and stored at -80°C.

### GST pull down

Recombinant GST proteins bound to glutathione agarose were incubated in a rotator with treated or naïve cell lysate 16h at 4°C. Beads were washed with NP40 buffer 6×300μl and recombinant GST and associated proteins were eluted from glutathione agarose beads with 70μl of 10mM glutathione in 50mM Tris-HCl pH 9.5.

### In-vitro degradation assay

Degradation of recombinant baculovirus expressed and purified p53 (kindly provided by Prof. C. Prives, Columbia University, New York) and IDP enriched lysate by purified 20S proteasomes was carried out in degradation buffer [100mM Tris-HCL pH 7.5, 150mM NaCl, 5mM MgCl_2_, 2mM DTT] at 37ºC for 1 hour and 3 hours respectively. The degradation reaction was stopped with the addition of Laemmli sample buffer to the samples. The samples were then heated at 95ºC for 5 min and fractionated by SDS-PAGE. Following electrophoresis, proteins were visualized by gel staining or transferred to cellulose nitrate membranes. Purified 20S proteasome was detected by immunoblotting with rabbit anti-PSMA4 antibody. Baculovirus-expressed and purified p53 was detected by immunobloting with mouse anti-human p53 (1801).

### Purified p53

Infection and purification of recombinant baculovirus-expressed human p53 from insect cells was done as described ^83^.

### Data analysis

Data analysis was preformed with a web-tool for plotting box plots (http://boxplot.tyerslab.com) and Microsoft Excel.

## Acknowledgments

We would like to thank Gad Asher for critical reading of the paper, Tsviya Olender for help with bioinformatics, Chaim Kahana and Keiji Tanaka for providing essential materials. This research was supported by the Israel Science Foundation (grant no. 1591/15) and the Kahn Family Research Center.

---

Supplementary table 1: **PSMA subunits interacting proteins.** We searched the IMEx data resource for identified protein-protein interactions identified in large-scale screens for each subunit. Submitted as an excel file.

**Supplementary 1:**
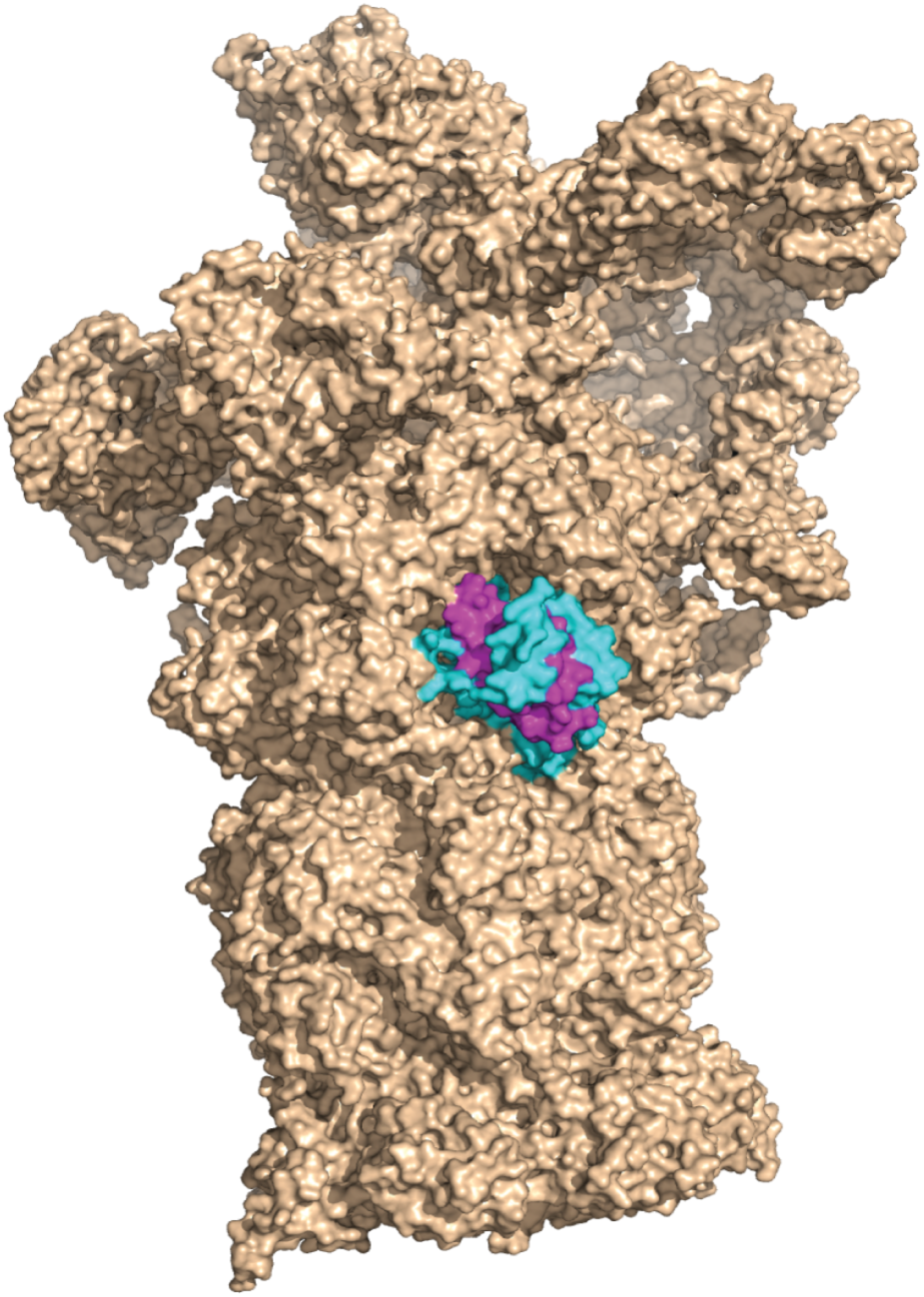
PSMA3 Ct is exposed in the 26S proteasome. Cryo-EM structure of the 26S proteasome marking PSMA3 Ct adapted from Huang *et al.*, 2016.

**Supplementary 2:**
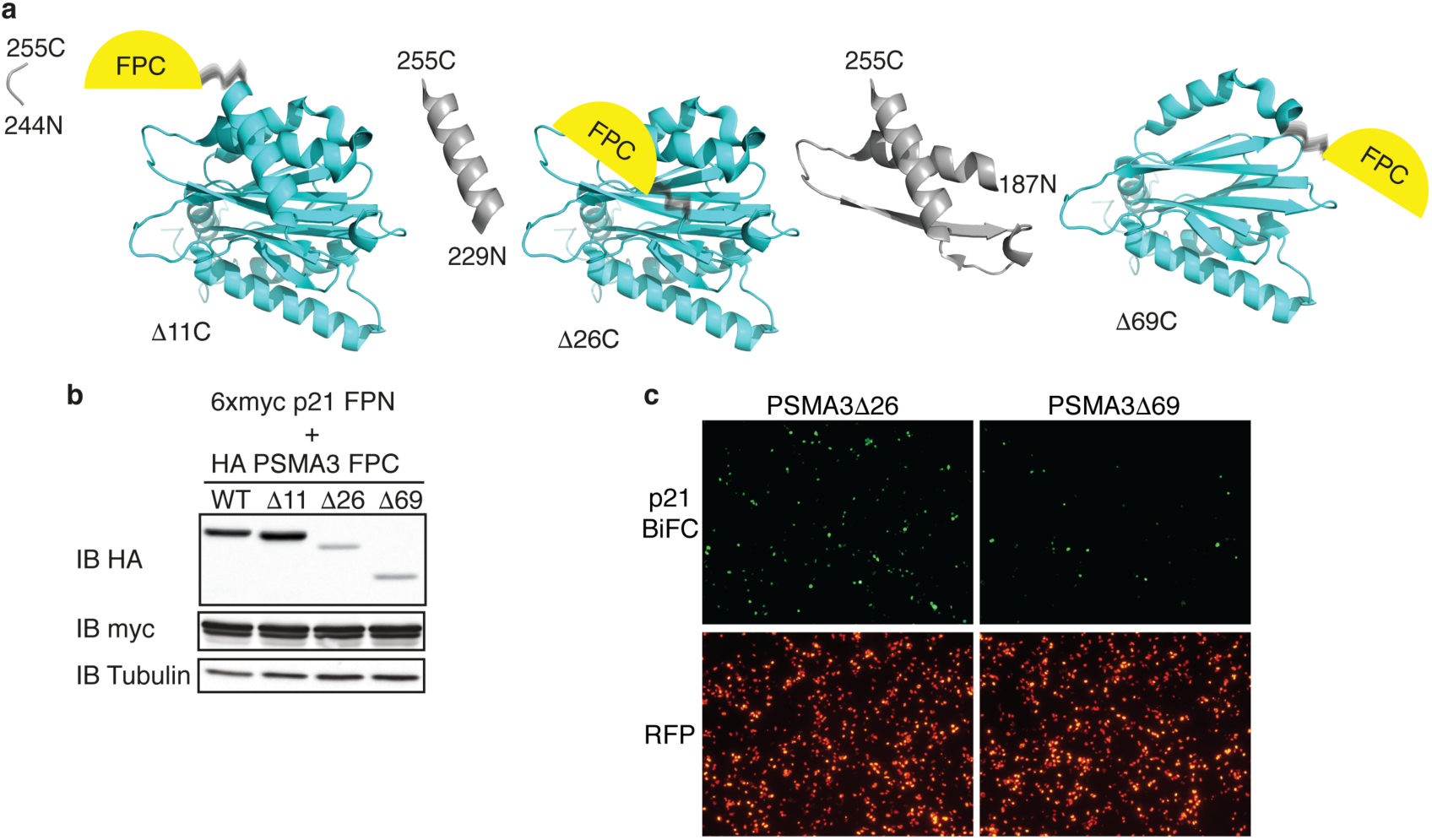
Identification by truncation mutagenesis of the PSMA3 region mediating the interaction with the p21. (a) Chimeric PSMA3 C-terminus deletion constructs used to identify the interaction region with p21. Truncated regions are colored in gray. Crystal structure adapted from Schrader *et al.*, 2016. (b-c) HEK293 cells were transiently transfected as indicated with 6xmyc p21 FPN, chimeric PSMA3 WT, deletion mutants and H2B RFP constructs. (b) Expression level of proteins and (c) successful BiFC were examined. Images were taken with a fluorescent microscope, 10x objective 72h after transfection.

**Supplementary 3:**
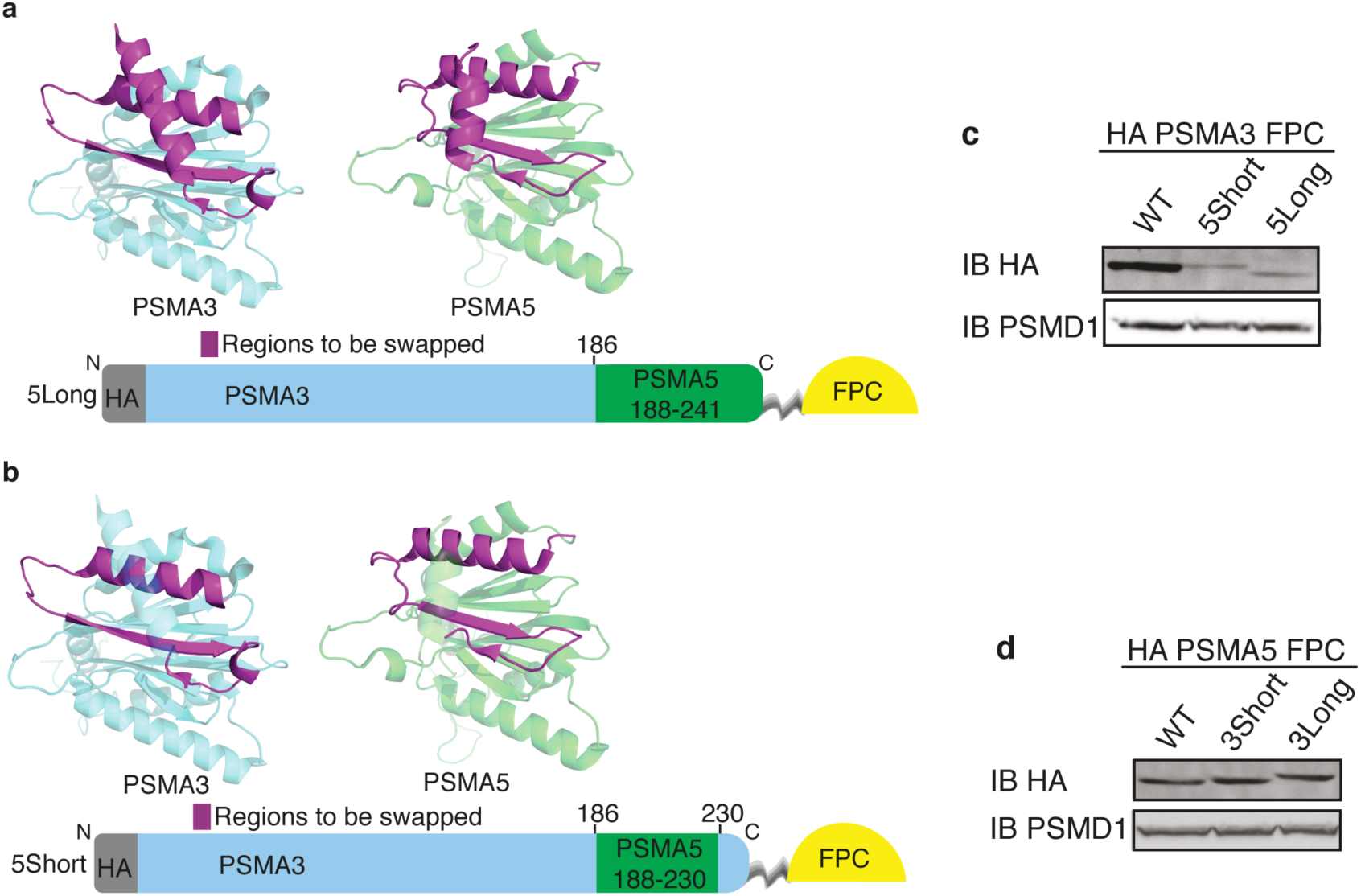
Expression of the PSMA3,5 chimeric constructs. (a-b) Illustration of mutant PSMA3 constructs used. Amino acid notations refer to PSMA3 WT. Crystal structures adapted from Schrader *et al.*, 2016. (c) Expression of PSMA3 swap proteins. Immunoblot (IB). (d) Expression of PSMA5 swap proteins. Immunoblot (IB).

**Supplementary 4:**
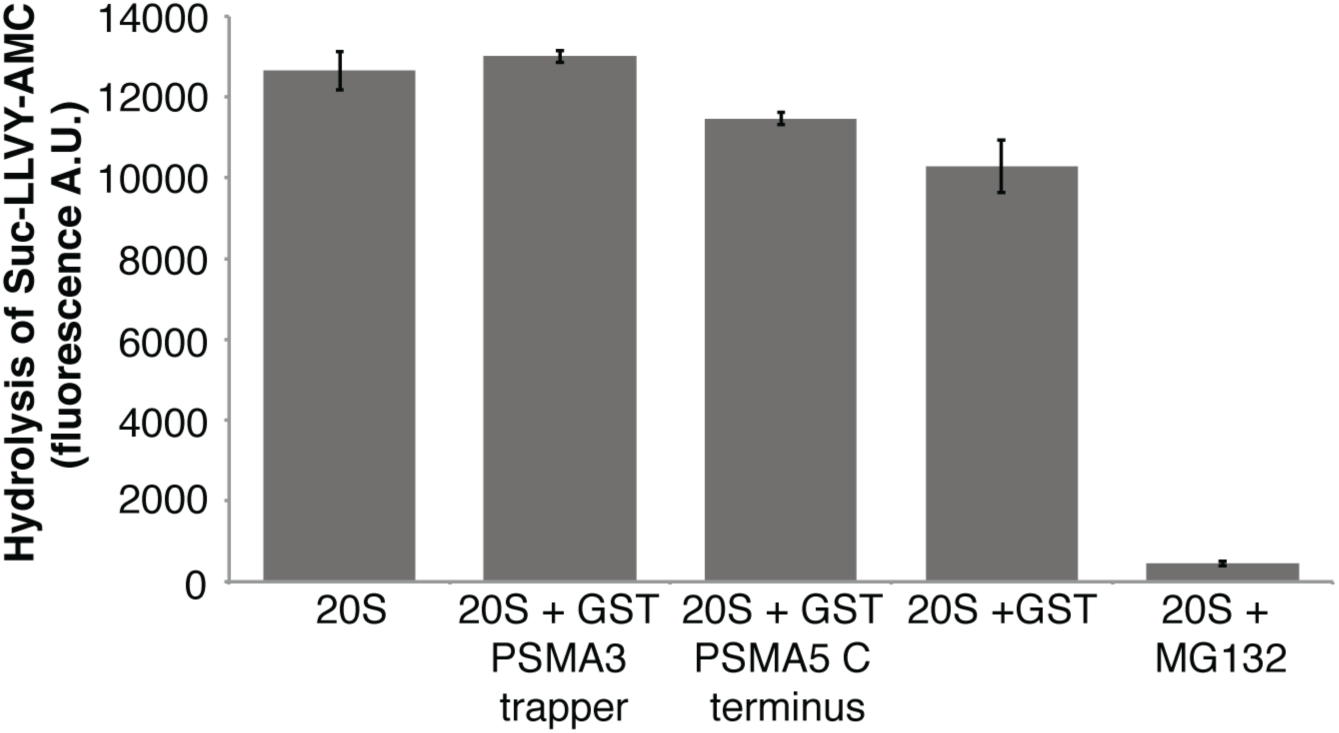
The PSMA3 trapper does not inhibit proteasome catalytic activity. Purified 20S proteasome was incubated 30 minutes at 37°C as indicated with purified GST, GST PSMA3 trapper, GST PSMA5 C-terminus and proteasome inhibitor MG132 in the presence of the chymotrypsin-like fluorogenic substrate Suc-LLVY-AMC. Standard deviation bars represent three independent experiments.

**Supplementary 5:**
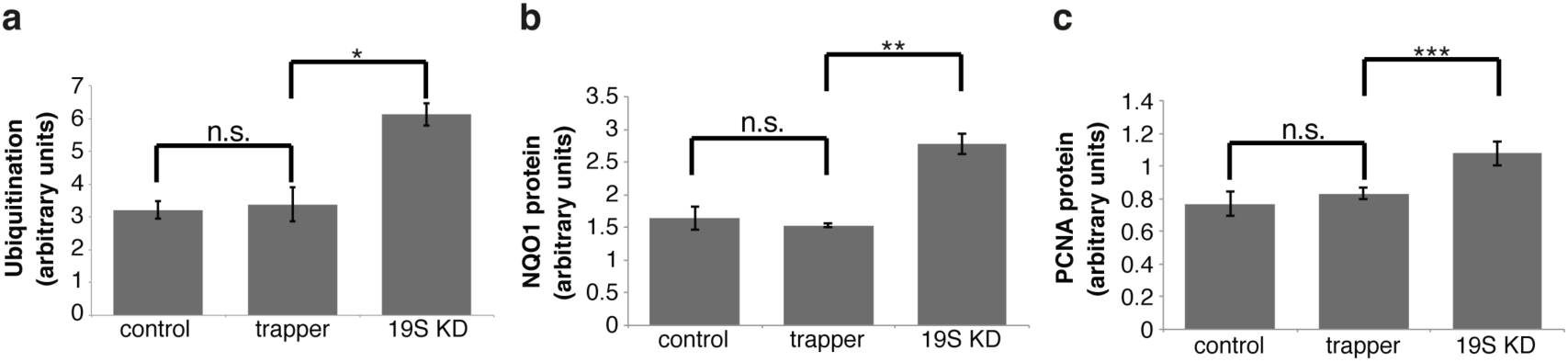
PSMA3 trapper does not inhibit ubiquitin dependent degradation. (a-c) HCT116 cells were infected with lentivirions based on the doxycycline-inducible vector pTRIPZ PSMD1 shRNA and selected with 2μg/ml puromycin. These cells were used for additional infection with lentivirions based on pLenti6 YFP or YFP fusions with PSMA3 trapper and selected with 10μg/ml blasticidin. To induce PSMD1 shRNA expression, the cells were treated with 1μg/ml doxycycline for 3 days and harvested for Western blot. PSMD1 knock down decreases 19S proteasome and inhibits ubiquitin dependent degradation. Western blots of (a) total ubiquitination (b) NQO1 protein and (c) PCNA proteins were quantified. *pValue=0.004 **pValue=0.00003 ***pValue=0.0007 using 2 sided student t test. n.s. is a non significant pValue.

